# Ascorbate and TET2 couple balanced hematopoiesis to disease tolerance

**DOI:** 10.64898/2026.10.27.762224

**Authors:** Tobias Wijshake, Joseph Rose, Ji Hyung Jun, Jacob Zielke, Li Li, Joseph S. Takahashi, Filipa Rijo-Ferreira, Michalis Agathocleous

## Abstract

Infections stimulate myelopoiesis but this comes at the cost of reduced production of other lineages. It is unclear if this imbalance is protective or damaging, and if factors that promote balanced hematopoiesis can impact infection pathology. Here we show that the ascorbate-TET axis limits excessive myelopoiesis, promotes balanced hematopoiesis, and is essential for surviving malaria. Hematopoietic-specific ascorbate depletion promoted myelopoiesis. The effects of ascorbate were mediated by TET2/TET3 activity in myeloid progenitors. Combined ascorbate and TET2 depletion markedly skewed hematopoiesis to the myeloid lineage and impaired erythropoiesis. Ascorbate is commonly depleted in infections, but the functional significance of this remains unclear. In the absence of ascorbate and TET2, mice infected with *Plasmodium* parasites could clear the pathogen, but had exacerbated myeloid and impaired erythroid response, anemia, and tissue damage and they succumbed to infection, indicating compromised disease tolerance. Therefore ascorbate and TET2 promote balanced hematopoiesis to impart tolerance to malaria.

## INTRODUCTION

Infection with the same microbe can kill some individuals even when most have mild or no disease, a phenomenon termed ‘the infection enigma’^1,2^. One cause is heritable host variation that affects pathogen load, such as protective mutations that block pathogen cellular entry or growth, and deleterious mutations that increase pathogen growth by causing immunodeficiencies^1,2^. A second cause may be non-heritable host variation that affects pathology without affecting pathogen load. Malaria is a classic example as protection from severe disease does not rely solely on restricting parasite load, suggesting the infected host averts severe pathology by tolerating the parasite^3^.

A balanced hematopoietic output is essential for appropriate immune and blood cell function. In infection, hematopoietic stem and progenitor cells (HSPCs) increase myelopoiesis to clear the pathogen, but at the cost of reducing production of other cell types, including lymphoid and erythroid cells. Emerging evidence suggests that reduced lymphopoiesis exacerbates the pathology of some infections by impairing antimicrobial immunity^4^. Whether imbalanced hematopoiesis affects the pathology of infection independently of affecting pathogen load has not been tested.

The metabolite ascorbate acts as a signal that controls hematopoiesis. Ascorbate levels are high in hematopoietic stem cells (HSCs) and decline with differentiation, and ascorbate depletion stimulates HSC function^5^. Ascorbate is taken up by hematopoietic cells through its dedicated transporter SLC23A2, and *Slc23a2* deletion stimulates HSC and multipotent progenitor (MPP) function^5,6^. Ascorbate promotes T cell, plasma cell, and erythroid development via effects at distinct differentiation steps^7–11^. These results raise the question of whether ascorbate has a coordinating role in hematopoietic output by suppressing the development of other lineages.

Most mammals synthesize ascorbate in the liver using the enzyme L-gulonolactone oxidase (GULO). Some species including humans lost *GULO* in evolution, turning ascorbate into an essential micronutrient better known as vitamin C^12,13^. As a result, low ascorbate levels are common in the population^14,15^. Low ascorbate is associated with a higher risk of severe respiratory infections^16^. Ascorbate stores are further depleted in several infections or inflammation^17–20^. Ascorbate depletion in *Gulo^−/−^*mice increases the pathology of influenza^21^, pneumonia^22^ and bacterial sepsis^23^. However ascorbate supplementation trials in patients with bacterial or viral infections showed mixed results, with some showing a benefit and others not^24–27^. In a recent phase 2 randomized trial, ascorbate supplementation of patients with low-risk myeloid malignancies or pre-malignancies cut mortality by half. The benefit was driven by prevention of severe infections through unknown mechanisms^28^. Supplemental ascorbate during infections is used widely in the population, and its role has been debated for decades^25,29^. Despite this, the mechanisms by which ascorbate operates in infections remain unclear.

## RESULTS

### Hematopoietic ascorbate depletion induces myeloid skewing

To understand the cell-intrinsic role of ascorbate in hematopoiesis, we depleted ascorbate in hematopoietic cells but not systemically by conditionally deleting the ascorbate transporter *Slc23a2* with *Vav1Cre*. Ascorbate was depleted in *Slc23a2^Δ/Δ^* bone marrow cells (**Figure 1A**), consistent with previous findings that hematopoietic *Slc23a2* deletion depletes ascorbate in hematopoietic cells but not the plasma^5,6^. *Vav1Cre;Slc23a2^Δ/Δ^* mice had a hypocellular bone marrow with reduced frequency and number of hematopoietic stem cells (HSCs), multipotent progenitors (MPPs) and hematopoietic progenitor cells (HPC-1) in the bone marrow, and increased frequency or number of common myeloid progenitors (CMPs) in the bone marrow and spleen and granulocyte monocyte progenitors (GMPs) in the spleen (**Figure 1B-C, Figure S1A-B**). *Vav1Cre;Slc23a2^Δ/Δ^* mice had increased frequencies or numbers of myeloid cells, particularly inflammatory monocytes in bone marrow, spleen and blood (**Figure 1D, Figure S1C-D**). Some lymphoid cells were reduced, and red blood cell and platelet counts were largely unaffected (**Figure S1C-F**).

**Figure 1.**
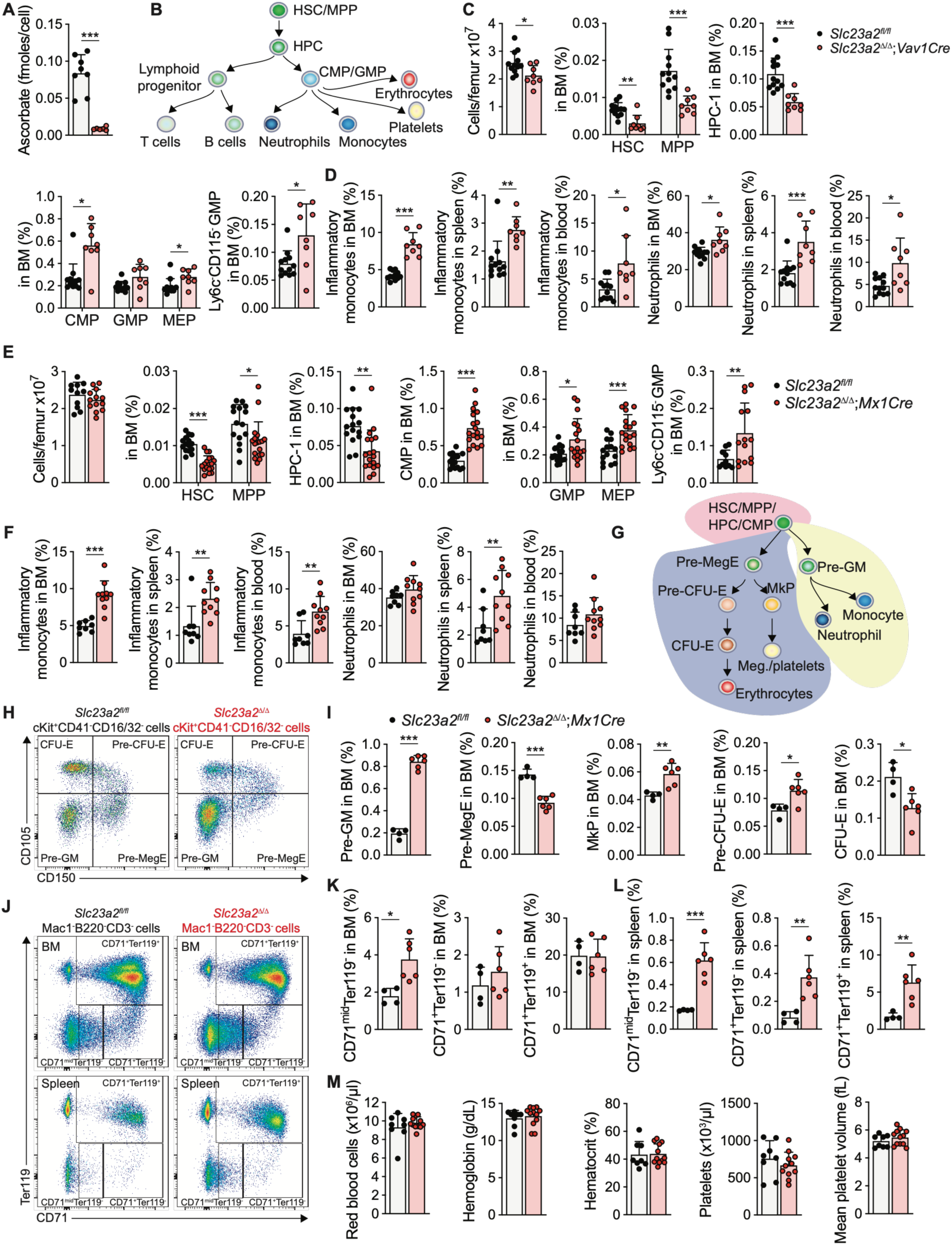
Hematopoietic ascorbate deficiency promotes myelopoiesis. (**A**) Intracellular ascorbate levels in bone marrow cells of *Vav1Cre;Slc23a2*^Δ/Δ^ and littermate control mice. (**B**) Schematic of hematopoietic stem cell differentiation. (**C-D**) Cellularity and frequencies of hematopoietic cells in the bone marrow of *Vav1Cre;Slc23a2*^Δ/Δ^ and littermate control mice at 8 weeks of age. (**E-F**) Cellularity and frequencies of hematopoietic cells in the bone marrow of wild-type littermate control and *Mx1Cre;Slc23a2*^Δ/Δ^ mice at 4-8 weeks after deletion with poly(I:C). (G) Schematic of myeloerythroid differentiation. (**H-I**) Flow plots and frequencies of erythroid, megakaryocytic and myeloid progenitors in the bone marrow of *Mx1Cre;Slc23a2*^Δ/Δ^ and littermate control mice. (**J-L**) Flow plots and frequencies of erythroid progenitors in the bone marrow and spleen. (**M**) Blood cell counts of wild-type and *Mx1Cre;Slc23a2*^Δ/Δ^ mice. *In all figures, data represent mean ± s.dev., individual dots in bar graphs represent biological replicates (mice), and statistical significance was assessed as described in the methods. *p < 0.05, **p < 0.01, ***p <0.001*.

Because *Vav1Cre* deletes starting from embryonic development we next asked if ascorbate is required in adult hematopoiesis by generating *Mx1Cre;Slc23a2^fl/fl^* mice in which deletion was induced with poly (I:C) injection. *Slc23a2* deletion in young adult mice had no effect on bone marrow cellularity and reduced bone marrow HSCs, MPPs, and HPC-1 consistent with prior work ^6^. *Slc23a2* deletion increased CMPs, GMPs, other restricted myeloid progenitors and inflammatory monocytes in the marrow, spleen, or blood (**Figure 1E-F, Figure S2A-D**). B cells were reduced and T cells were unaffected (**Figure S2C-E**). Therefore, ascorbate deficiency in adult hematopoiesis induces myelomonocytic skewing.

We then examined if the increase in restricted myeloid progenitors reflected an increase in specific subpopulations. Restricted myeloid progenitors contain cells with megakaryocyte-erythroid or monocytic-granulocytic potential^30,31^ (**Figure 1G**). *Slc23a2* deletion increased pre-granulocyte-macrophage progenitor (Pre-GM) cells and reduced pre-megakaryocyte-erythrocyte progenitors (Pre-MegE). It increased megakaryocyte progenitors (MkP) and pre-colony-forming-unit-erythroid progenitors (Pre-CFU-E), and decreased CFU-Es in the bone marrow (**Figure 1H-I, Figure S2F**). Late-stage erythropoiesis was largely unimpaired in bone marrow and increased in the spleen, and *Slc23a2^Δ/Δ^* mice maintained red blood cell and platelets counts **(Figure 1J-M, Figure S2G-H**). Therefore, *Slc23a2^Δ/Δ^* myelomonocytic expansion was accompanied by mild impairments in megakaryocyte-erythroid differentiation that did not result in overt red blood cell or platelet deficits at baseline.

### Ascorbate acts in oligopotent and restricted myeloid progenitors to limit myelopoiesis

To determine whether ascorbate suppresses myeloid cell numbers by acting in mature myeloid cells or upstream progenitors, we generated *LysMCre;Slc23a2^Δ/Δ^* mice. *LysMCre* deletes in most granulocytes and monocytes but in only ∼10% of HSCs and restricted progenitors^32^. In contrast to pan-hematopoietic deletion, *LysMCre-*driven *Slc23a2* deletion caused no changes in cell number or HSC and progenitor frequency in the bone marrow (**Figure 2A, Figure S3A-B**). Inflammatory monocytes and neutrophils slightly increased in the bone marrow, but not the spleen or blood (**Figure 2B, Figure S3C-D**). These increases were much smaller in magnitude than those in *Mx1Cre;Slc23a2^Δ/Δ^* or *Vav1Cre;Slc23a2^Δ/Δ^* mice (**Figure 1D**, **Figure 1F**), suggesting that most or all the effect of *Slc23a2* deletion in increasing myeloid cell number was mediated by actions upstream of mature myeloid cells.

**Figure 2.**
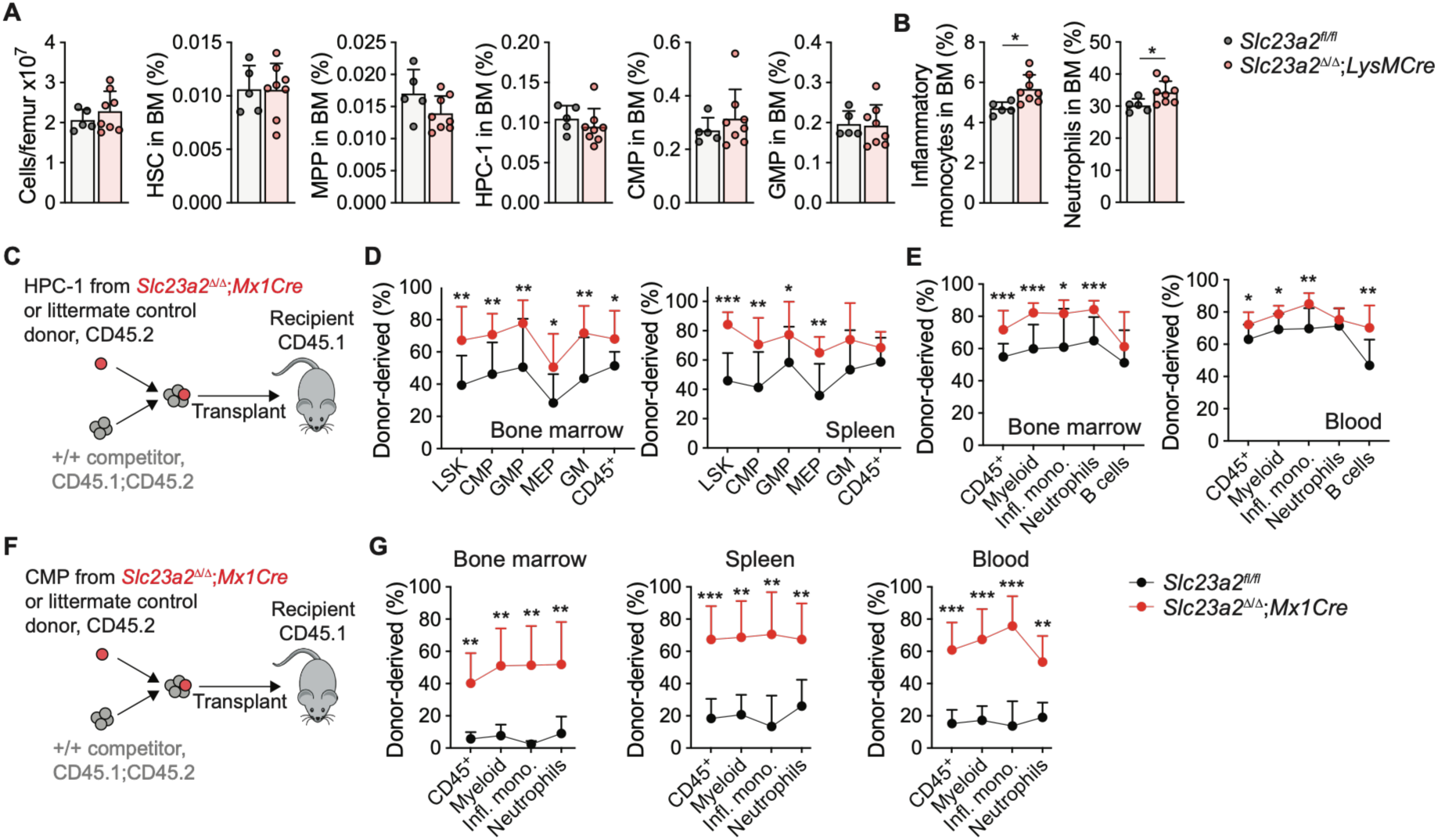
Ascorbate inhibits myelopoiesis by acting in oligopotent and restricted myeloid progenitors. (**A-B**) Bone marrow cellularity and the frequencies of hematopoietic stem cells, progenitors, and mature cells in bone marrow of *LysMCre;Slc23a2*^Δ/Δ^ and littermate control mice at 8 weeks of age. (**C-E**) Competitive transplant to assess ascorbate-deficient HPC-1 reconstitution function. 1,000 HPC-1 (CD45.2) from *Mx1Cre;Slc23a2*^Δ/Δ^ or littermate control mice were mixed with 400,000 competitor bone marrow (CD45.1;CD45.2) cells from wild-type mice and transplanted to lethally irradiated recipient (CD45.1) mice (*n* = 13 recipient mice/genotype from 3 donors/genotype). (**D-E**) Donor chimerism of myeloid progenitors in bone marrow and spleen, and mature cells in bone marrow and blood 14-15 days after transplant. (**F-G**) Competitive transplantation to assess ascorbate-deficient CMP reconstitution function. 5,000 CMPs (CD45.2) from *Mx1Cre;Slc23a2*^Δ/Δ^ or littermate control mice were mixed with 400,000 competitor bone marrow (CD45.1;CD45.2) cells from wild-type mice and transplanted to lethally irradiated recipient (CD45.1) mice (*n* = 13-14 recipient mice/genotype from 3 donors/genotype). (**G**) Donor chimerism of mature cells in bone marrow, spleen and blood 12-14 days after transplant.

We next tested if ascorbate suppresses myeloid output by acting in upstream progenitors. Ascorbate acts within HSCs and MPPs to suppress self-renewal^5,6^. To investigate if ascorbate acts within oligopotent progenitors to suppress myelopoiesis, we transplanted 1,000 purified CD150^−^CD48^−^Lin^−^Sca1^+^Kit^+^ HPC-1 cells^33^ from *Mx1Cre;Slc23a2^Δ/Δ^* or wild-type mice with wild-type bone marrow competitor cells in irradiated recipient mice (**Figure 2C**). 2 weeks later, *Slc23a2* deletion increased reconstitution of progenitors and mature myeloid cells (**Figure 2D-E**). *Slc23a2^Δ/Δ^* HPC-1 B cell production was unchanged in the marrow and slightly increased in the blood (**Figure 2E**). Together with data that *Slc23a2^Δ/Δ^*HSCs/MPPs produce more B cells than wild-type cells after competitive transplantation^5,6^, this suggested that the decreased B cell frequency in *Slc23a2^Δ/Δ^* mice was caused by non-cell-autonomous mechanisms. To investigate if ascorbate suppresses myelopoiesis by acting within restricted myeloid progenitors, we transplanted 5,000 purified *Slc23a2*^Δ/Δ^ CMPs with wild-type competitors in irradiated recipient mice (**Figure 2F**). *Slc23a2* deletion dramatically increased CMP-derived myeloid reconstitution in bone marrow, spleen and blood (**Figure 2G**). Therefore, ascorbate action within oligopotent and restricted myeloid progenitors suppresses myelopoiesis.

### Ascorbate deficiency collaborates with *Tet2* deficiency to stimulate myelopoiesis and impair erythropoiesis

Ascorbate acts as a cofactor that promotes the activity of the TET1-3 family of α-ketoglutarate-dependent dioxygenases that oxidize 5-methylcytosine on nucleic acids^34^. Ascorbate depletion in *Gulo*^−/−^mice reduces activity of TET2 in HSCs and of multiple TET enzymes in myeloid progenitors or myeloid leukemia^5^. Somatic *TET2* mutations in the blood system are prevalent in humans and cause clonal HSC expansion and myeloid neoplasms^35^. *Tet2* deletion in mice increases HSC self-renewal and myelopoiesis^36–40^. To test if *TET2* deletion synergizes with ascorbate depletion to exacerbate myeloid skewing, we generated *Mx1Cre;Slc23a2^Δ/Δ^;Tet2^Δ/Δ^* mice. Combined loss of *Slc23a2* and *Tet2* in adult mice caused a two to four-fold increase in myeloid progenitors in the bone marrow and spleen (**Figure 3A, Figure S4A-B**). These effects were stronger than those of single *Slc23a2* deletion (**Figure 1E**) or *Tet2* deletion, which causes a ∼30% increase in CMP frequency and no change in GMP frequency^5^. Consequently, inflammatory monocytes and neutrophils accumulated to a greater extent in *Slc23a2^Δ/Δ^;Tet2^Δ/Δ^* than in *Slc23a2^Δ/Δ^* mice or *Tet2^Δ/Δ^* mice (**Figure 3B**). Myeloid expansion was accompanied by a contraction of other lineages (**Figure S4C-F**). *Slc23a2*;*Tet2* deletion increased the frequency of pre-CFU-E, CD71^mid^Ter119^−^ and CD71^+^Ter119^−^erythroid progenitors yet depleted CD71^+^Ter119^+^ erythroblasts in the bone marrow, suggesting an erythropoietic arrest (**Figure 3C-F)**. *Slc23a2*;*Tet2* deletion increased megakaryocyte progenitors and spleen erythropoiesis (**Figure 3D-E, G**) yet this was accompanied by mild anemia and thrombocytopenia (**Figure 3H**), consistent with the effects on progenitors reflecting compensatory hematopoiesis. Therefore, ascorbate deficiency collaborates with TET2 deficiency to enhance myelopoiesis and impair erythropoiesis.

**Figure 3.**
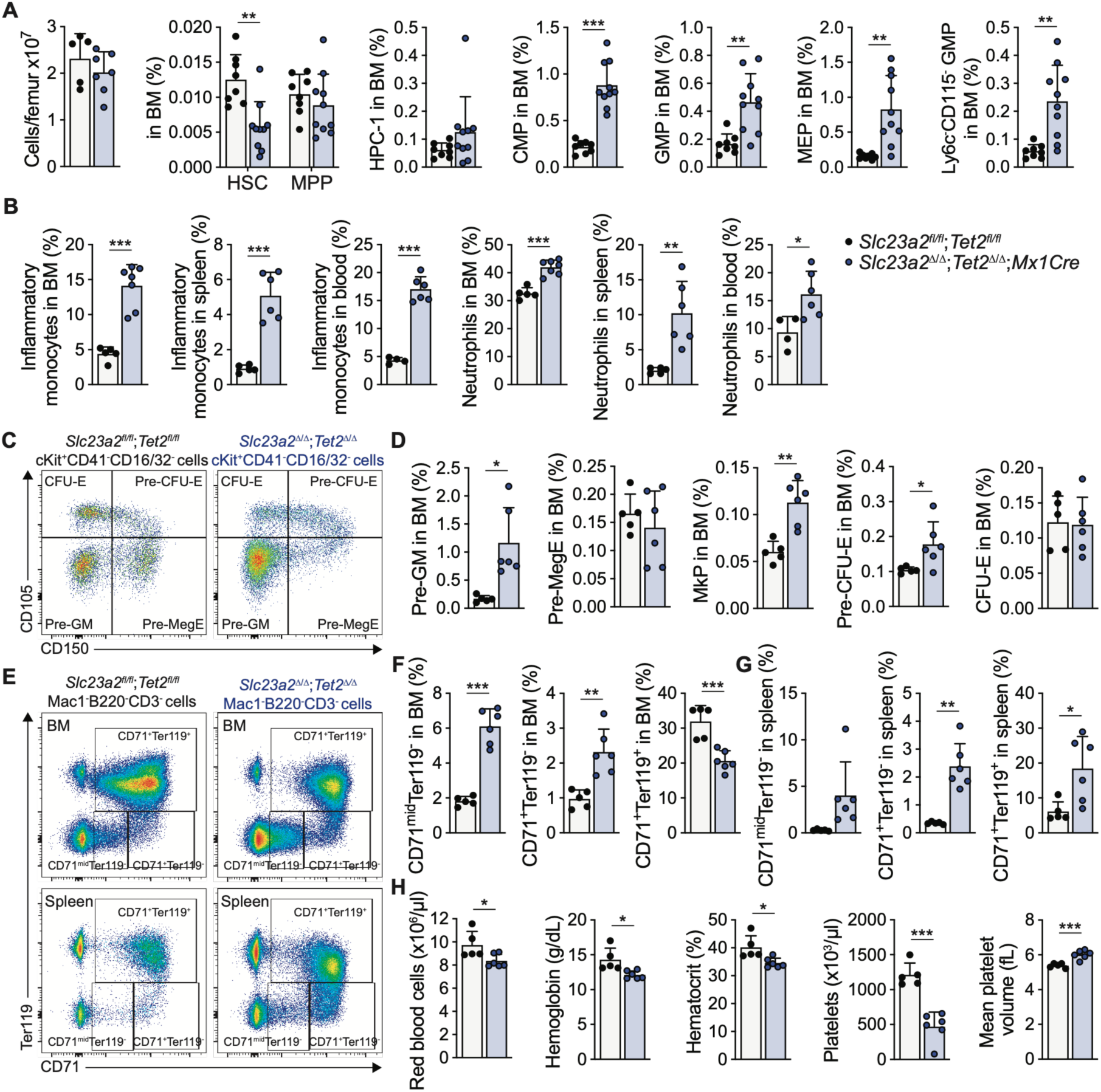
Ascorbate deficiency and TET2 deficiency collaborate to promote myelopoiesis and impair erythropoiesis. (**A-B**) Cellularity and the frequencies of hematopoietic stem cells, progenitors (**A**), and mature myeloid cells (**B**) in the bone marrow of *Mx1Cre;Slc23a2*^Δ/Δ^*;Tet2*^Δ/Δ^ or littermate control mice 4-8 weeks after *Mx1Cre* induction with poly(I:C). (**C-D**) Flow plots and frequencies of erythroid, megakaryocytic and myeloid progenitors in bone marrow. (**E-G**) Flow plots and frequencies of erythroid progenitors in bone marrow and spleen of the indicated mice. (H) Blood cell counts of wild-type and *Mx1Cre;Slc23a2*^Δ/Δ^*;Tet2*^Δ/Δ^ mice.

### Ascorbate restrains myelopoiesis by stimulating TET2/TET3 activity

To test if ascorbate deficiency increased myelopoiesis by inhibiting TET2 and TET3, we generated *Mx1Cre;Tet2^Δ/Δ^, Mx1Cre;Tet3^Δ/Δ^*, *Mx1Cre;Tet3^Δ/Δ^;Slc23a2^Δ/Δ^,* and *Mx1Cre;Tet2^Δ/Δ^;Tet3^Δ/Δ^* mice. We measured levels of the TET product 5-hydroxymethylcytosine (5hmC) on hydrolyzed DNA from purified HSC and progenitor cells 4-7 weeks after deletion (**Figure 4A**). HSC/MPPs had the highest 5hmC levels among analyzed cell types, consistent with prior data^5^ (**Figure 4B**). *Tet3* deletion had no effect on total TET activity in any cell type. *Tet2* deletion mildly reduced TET activity in HSPCs and strongly reduced it in restricted myeloid progenitors. Combined *Tet2;Tet3* deletion decreased TET activity more than individual *Tet2* or *Tet3* deletion suggesting TET2 and TET3 compensate for each other in all cell types. *Slc23a2* deletion decreased TET activity more than individual *Tet2* or *Tet3* deletion, suggesting it inhibits multiple TET enzymes. Combined *Slc23a2;Tet2* deletion decreased TET activity to a similar extent as combined *Tet2;Tet3* deletion (**Figure 4B**). Despite highly decreased 5hmC levels, total 5mC levels in all genotypes were unaffected, consistent with 5hmC being a stable mark in HSPCs as shown in other tissues^41^.

**Figure 4.**
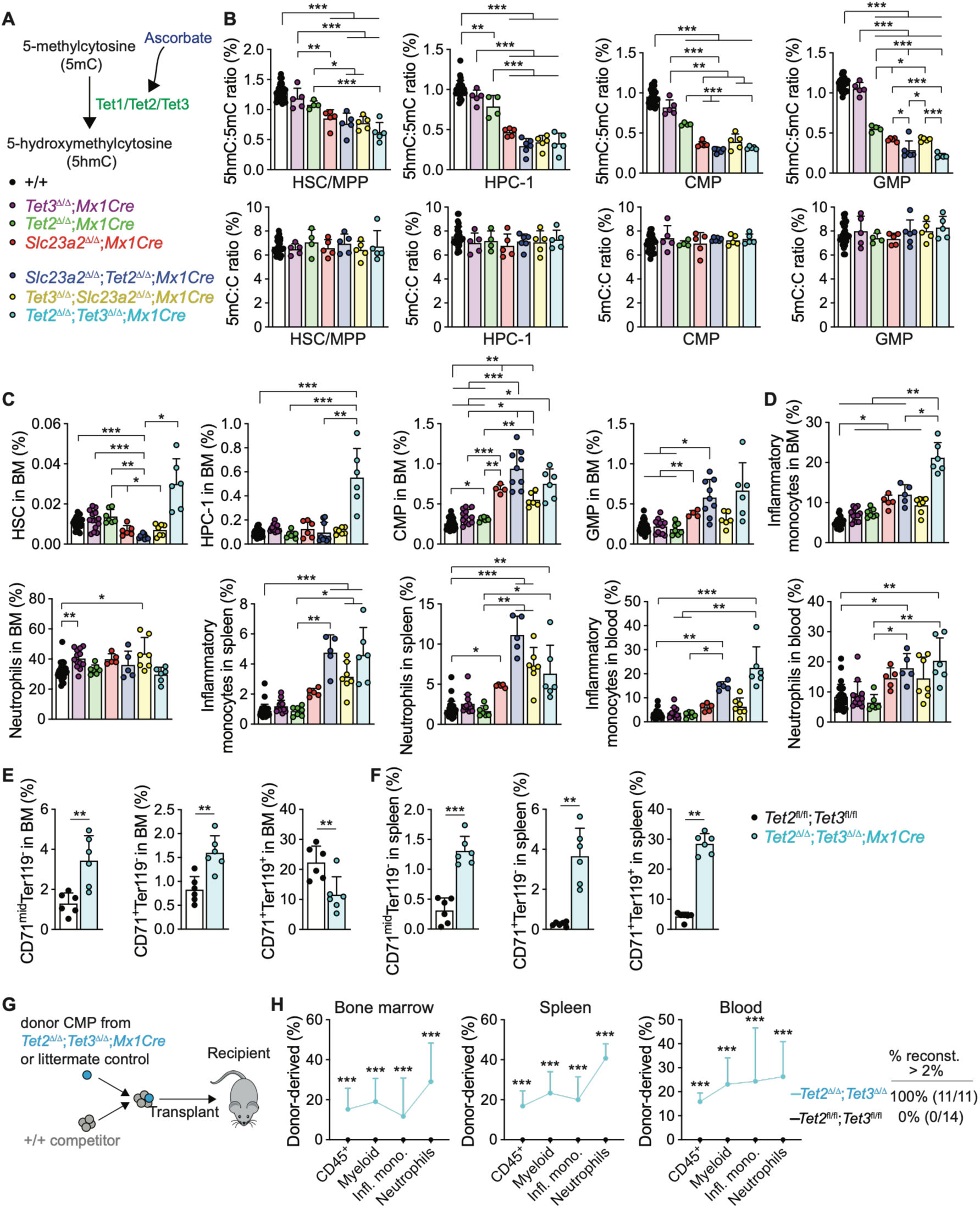
The effects of ascorbate deficiency on myelopoiesis are phenocopied by *Tet2;Tet3* deficiency. (**A**) Ascorbate enhances Tet catalytic activity and promotes the conversion of 5-methylcytosine (5mC) to 5-hydroxymethylcytosine (5hmC) on DNA. (**B**) 5hmC:5mC and 5mC:C ratio on DNA of sorted HSCs and progenitors from mice with the indicated genotypes. +/+ refers to littermate control mice carrying corresponding floxed alleles. (**C-D**) Frequencies of HSPCs and myeloid cells in from mice with the indicated genotypes. (**E-F**) Frequencies of erythroid progenitors in bone marrow and spleen. (**G-H**) Competitive transplantation to assess *Tet2;Tet3*-deficient CMP reconstitution function. 5,000 CMPs (CD45.2) from *Mx1Cre;Tet2*^Δ/Δ^*;Tet3*^Δ/Δ^ or littermate control mice were mixed with 400,000 competitor bone marrow (CD45.1;CD45.2) cells from wild-type mice and transplanted to lethally irradiated recipient (CD45.1) mice (*n* = 11-14 recipient mice/genotype from 3 donors/genotype). (**H**) Donor chimerism of mature cells in bone marrow, spleen and blood 12 days after CMP transplant.

We tested if the effects of ascorbate deficiency on myelopoiesis were phenocopied by reduced TET2 and TET3 activity by analyzing single and compound mutants. *Tet3* deletion alone had minimal effects on HSPC frequencies. *Tet2;Tet3* deletion caused synergistic effects on most hematopoietic parameters compared to single knockouts, suggesting TET2 and TET3 functionally compensate for each other, consistent with the 5hmC results. *Tet3;Slc23a2* deletion had similar effects on 5hmC levels and hematopoiesis as *Slc23a2* deletion alone suggesting that *Slc23a2-*deficient cells have minimal or absent TET3 activity. *Tet2^Δ/Δ^;Tet3^Δ/Δ^* mice had increased CMP, GMP, inflammatory monocytes, and neutrophils, similar to *Slc23a2^Δ/Δ^;Tet2^Δ/Δ^* mice, and increased HSC and HPC-1, in contrast to *Slc23a2^Δ/Δ^;Tet2^Δ/Δ^* mice. *Slc23a2^Δ/Δ^;Tet2^Δ/Δ^* and *Tet2^Δ/Δ^;Tet3^Δ/Δ^* mice both had expanded myeloid cells in the blood and spleen, and expanded spleen HSPCs (**Figure 4C-D, Figure S4G**). Together with the 5hmC results, these results suggested that *Slc23a2* deletion promoted myelopoiesis mainly via TET2/TET3 inhibition, and decreased HSC and HPC-1 frequencies via other mechanisms. The increased reconstitution ability of *Slc23a2^Δ/Δ^* HSCs^5,6^ and HPC-1 (**Figure 2D-E**) suggests that the reduced *Slc23a2^Δ/Δ^* HSC and HPC-1 frequency is caused by non-cell autonomous mechanisms. Similar to *Slc23a2;Tet2* deletion, *Tet2;Tet3* deletion reduced CD71^+^Ter119^+^ erythroblasts in the bone marrow and increased spleen erythropoiesis (**Figure 4E-F**). To test if *Tet2;Tet3* deletion cell-autonomously promotes myelopoiesis, we transplanted purified CMP cells into irradiated recipients (**Figure 4G**). *Tet2^Δ/Δ^;Tet3^Δ/Δ^* CMPs produced significantly more myeloid cells than wild-type CMPs (**Figure 4H**). The similar biochemical, phenotypic, and functional consequences of *Slc23a2* deletion and *Tet2;Tet3* deletion suggest that ascorbate suppresses CMP-derived myelopoiesis mainly by promoting TET2/TET3 activity.

### Ascorbate and Tet2 loss exacerbate the myeloid response and inhibit the erythroid response to *Plasmodium chabaudi* infection

To determine the physiological importance of ascorbate-TET control of myelopoiesis and erythropoiesis, we used malaria infection, which demands a tightly coordinated myeloerythroid response. Malaria triggers emergency myelopoiesis that helps clear the pathogen^42,43^, but uncontrolled myeloid inflammation damages tissues^44^. Erythropoiesis is required in malaria to replace infected and destroyed erythrocytes, and erythropoiesis inhibition causes severe anemia^45,46^. Therefore, we tested if ascorbate and TET activity are required for a balanced myeloerythroid output in malaria. Malaria depletes ascorbate in patients^47^. While ascorbate depletion in *Gulo^−/−^*mice does not affect survival in an acute, fatal challenge with *Plasmodium berghei*^48^, most human malaria infections are non-lethal. We therefore infected mice with *Plasmodium chabaudi AS,* which causes a resolving non-lethal rodent malaria^46^, allowing us to test the role of the ascorbate-TET axis in hematopoiesis and pathology.

*Mx1Cre;Slc23a2^Δ/Δ^* mice infected with *P. chabaudi* showed a typical expansion and resolution of parasitized erythrocytes, with minor differences in clearance kinetics compared to wild-type mice, suggesting ascorbate in hematopoietic lineage cells is not required for the acute immune response against the pathogen (**Figure 5A-B**). However, the elevated inflammatory monocytes and neutrophils observed in *P. chabaudi*-infected wild-type mice were further increased by *Slc23a2* deletion. (**Figure 5C**). Infected *Vav1Cre;Slc23a2*^Δ/Δ^ mice showed similar phenotypes as *Mx1Cre;Slc23a2^Δ/Δ^* mice, including normal acute clearance of the parasite from the blood but minor recrudescence at 2-3 weeks post-infection and a large increase in blood myeloid cells compared to infected wild-type mice (**Figure S5A-C**). Infected *Mx1Cre;Tet2^Δ/Δ^* mice showed no changes in parasitemia or blood myeloid cells compared to infected controls indicating *Tet2* is not normally required for the response to infection (**Figure S5D-F**). Infected *Slc23a2^Δ/Δ^;Tet2^Δ/Δ^* mice had increased peak parasitemia compared to controls, mostly cleared the parasite from the blood by 2 weeks, and showed a mild recrudescence at 3 weeks, similar to *Slc23a2* deletion (**Figure 5D-E**) and a marked, sustained expansion of inflammatory monocytes (**Figure 5F**).

**Figure 5.**
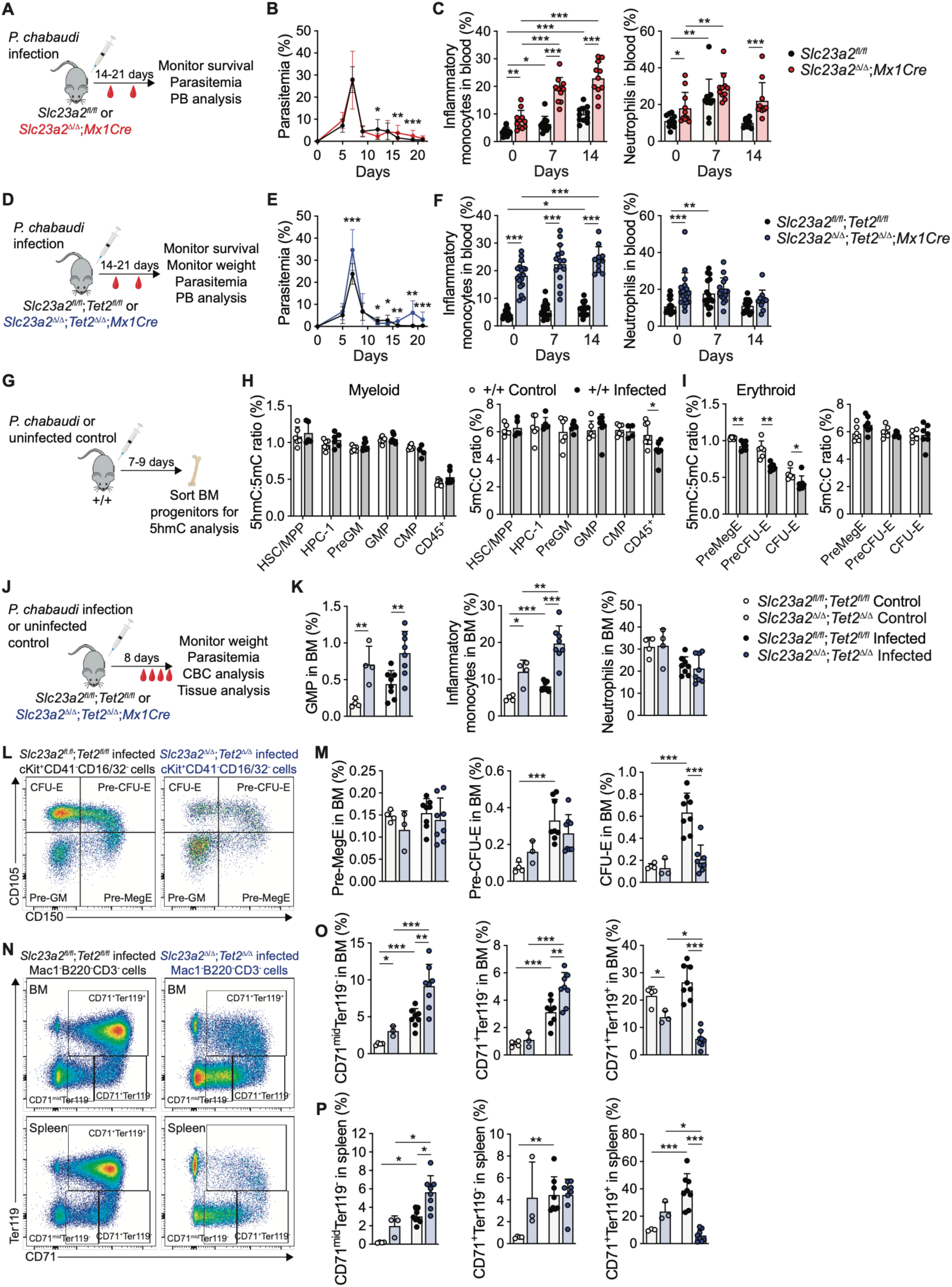
Combined ascorbate and TET2 deficiency cause imbalanced hematopoiesis in *Plasmodium chabaudi* infection. (**A-C**) *P. chabaudi* infection in *Mx1Cre;Slc23a2*^Δ/Δ^ mice or littermate controls, with parasitemia measured as % of infected erythrocytes, and peripheral blood analysis. (**D-F**) Parasitemia and blood analysis after *P. chabaudi* infection in *Mx1Cre;Slc23a2*^Δ/Δ^*;Tet2*^Δ/Δ^ mice or littermate controls. (**G-I**) Effect of *P. chabaudi* infection on 5hmC and 5mC levels on the DNA of purified HSCs and progenitors. (**J-P**) Hematopoietic analysis of wild-type and *Mx1Cre;Slc23a2*^Δ/Δ^*;Tet2*^Δ/Δ^ mice at the peak of *P. chabaudi* infection: (**K**) myeloid progenitors and mature cells; (**L-M**) flow plots and frequencies of megakaryocyte and erythroid progenitors; (**N-P**) flow plots and frequencies of late-stage erythroid progenitors.

To assess if *Plasmodium* infection impacts TET activity, we measured 5hmC levels in purified HSPCs from *P. chabaudi-*infected mice (**Figure 5G**). 5hmC levels were unchanged in HSCs and myeloid progenitors and decreased in erythroid lineage progenitors in infected mice, suggesting that infection limits TET activity in erythropoiesis (**Figure 5H-I**). At the peak of infection, *Slc23a2^Δ/Δ^;Tet2^Δ/Δ^* mice had increased bone marrow GMP and inflammatory monocyte frequency indicating increased myelopoiesis (**Figure 5J-K**). Infection markedly increased bone marrow CFU-E in wild-type mice but this was blocked in *Slc23a2^Δ/Δ^;Tet2^Δ/Δ^* mice (**Figure 5L-M**). Infected *Slc23a2^Δ/Δ^;Tet2^Δ/Δ^* mice also had blocked production of marrow CD71^+^Ter119^+^ erythroblasts (**Figure 5N-O**). Infection stimulated spleen erythropoiesis, but *Slc23a2;Tet2* deletion blunted this by blocking production of splenic erythroblasts (**Figure 5N-P**). Therefore hematopoietic *Slc23a2;Tet2* deficiency causes severe myeloerythroid imbalance in malaria by promoting the myelopoietic and inhibiting the erythropoietic response.

### Ascorbate and Tet2 loss impair tolerance to malaria

Imbalanced myeloerythroid hematopoiesis may have protective effects through parasite clearance or detrimental effects through inflammatory damage or anemia. Peak parasitemia increased in *Slc23a2^Δ/Δ^;Tet2^Δ/Δ^* mice (**Figure 5E**), suggesting the myeloid expansion did not promote parasite clearance. We thus tested if *Slc23a2^Δ/Δ^;Tet2^Δ/Δ^* mice had more severe malarial pathology. *Slc23a2^Δ/Δ^;Tet2^Δ/Δ^* mice exhibited severe morbidity. They had increased weight loss, an indicator of poor health in this model^49^ (**Figure 6A-B**). Consistent with blocked erythropoiesis, these mice had exacerbated anemia but not exacerbated thrombocytopenia (**Figure 6C-D**). *Slc23a2*^Δ/Δ^*;Tet2*^Δ/Δ^ mice had elevated inflammatory monocytes or neutrophils in tissues at baseline and malaria caused an additional increase in myeloid cells in tissues, including blood and liver, indicating increased inflammation (**Figure 6E, Figure S6A-B**). Infected *Slc23a2*^Δ/Δ^*;Tet2*^Δ/Δ^ mice had exacerbated liver damage indicated by alanine aminotransferase (ALT) and aspartate aminotransferase (AST)^44,50^, and hypoglycemia, a common malarial pathology (**Figure 6F-G**). Remarkably, while *P. chabaudi* infection resolved in wild-type control mice, 29% of *Slc23a2*^Δ/Δ^*;Tet2*^Δ/Δ^ mice succumbed to the disease (**Figure 6H**). Therefore, hematopoietic loss of ascorbate and TET2 increased the severity of malaria pathology and, in a subset of mice, transformed a resolving infection to a lethal disease.

**Figure 6.**
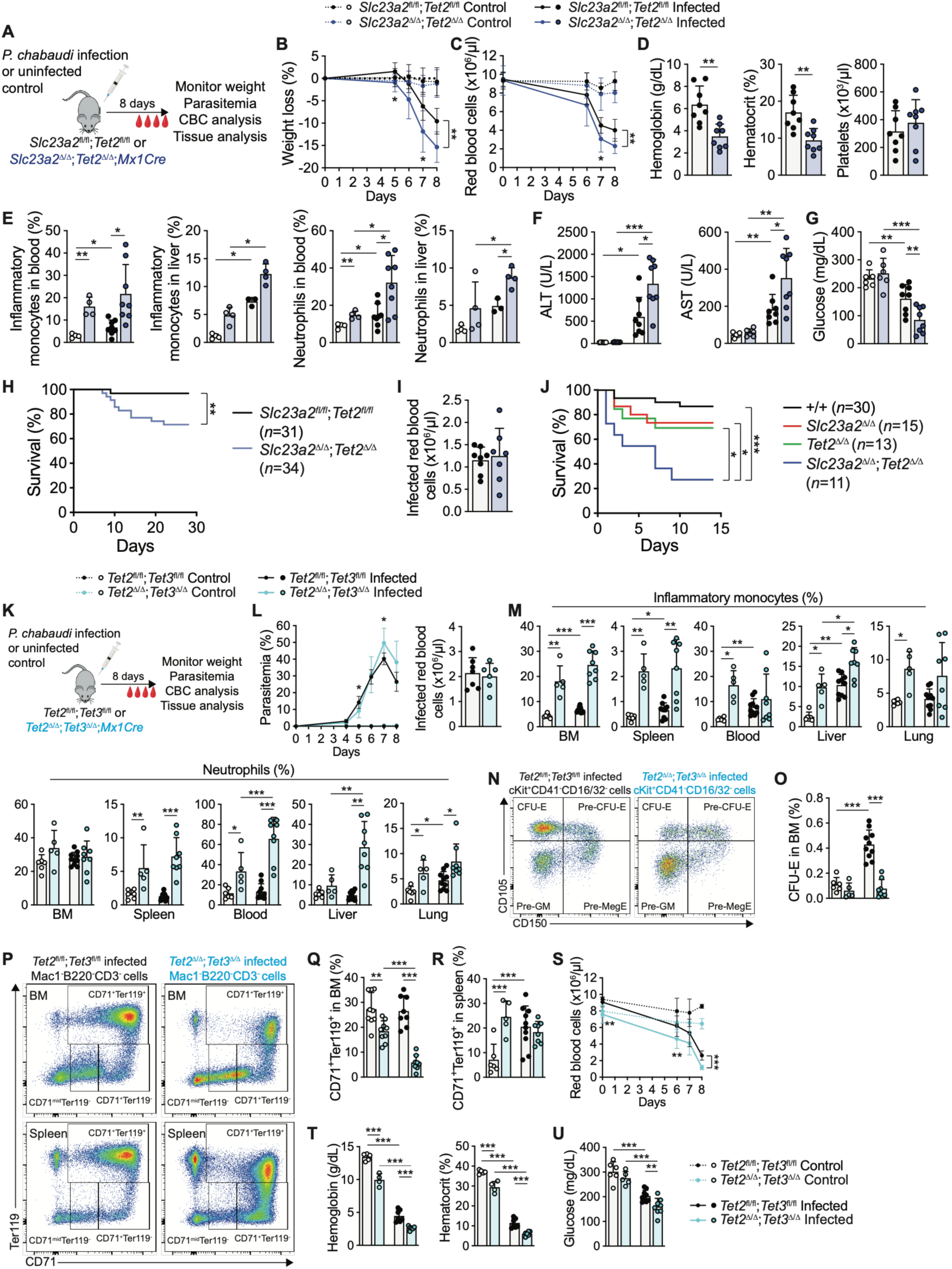
Combined ascorbate and Tet2 deficiency impair tolerance to *Plasmodium chabaudi* infection. (**A-G**) Measurement of malarial pathologies in *P. chabaudi*-infected wild-type and *Mx1Cre;Slc23a2*^Δ/Δ^*;Tet2*^Δ/Δ^ mice: (**B**) weight loss, (**C-D**) anemia, (**E**) frequency of myeloid cells in blood and liver, (**F-G**) liver damage and hypoglycemia, and (**H**) survival. (**I**) Number of infected erythrocytes at the peak of infection. (**J**) Survival of mice from the indicated genotypes after intraperitoneal injection of LPS (2.5mg/kg). +/+ refers to littermate control mice carrying the corresponding floxed alleles. (**K-R**) Analysis in wild-type and *Mx1Cre;Tet2*^Δ/Δ^*;Tet3*^Δ/Δ^ mice 8 days after infection. (**L**) Parasitemia and number of infected erythrocytes at peak of infection. (**M**) Frequencies of myeloid cells in the indicated tissues. (**N-O**) Flow plots and frequency of CFU-E in bone marrow. (**P-R**) Flow plots and frequencies of late-stage erythroid progenitors in bone marrow and spleen. (**S-U**) Anemia and hypoglycemia in infected wild-type and *Mx1Cre;Tet2*^Δ/Δ^*;Tet3*^Δ/Δ^ mice.

The decreased erythropoiesis in *Slc23a2*^Δ/Δ^*;Tet2*^Δ/Δ^ mice suggested that the increased frequency of parasitized erythrocytes may be a consequence of reduced erythrocyte production rather than increased parasite proliferation. The number of parasitized erythrocytes at the peak of parasitemia did not significantly differ in *Slc23a2*^Δ/Δ^*;Tet2*^Δ/Δ^ compared to wild-type mice (**Figure 6I**), suggesting increased parasite frequency in infected *Slc23a2*^Δ/Δ^*;Tet2*^Δ/Δ^ mice was due to the increased anemia caused by blocked erythropoiesis. The equal parasite load between wild-type and *Slc23a2*^Δ/Δ^*;Tet2*^Δ/Δ^ mice despite the large increase in disease severity and mortality in *Slc23a2*^Δ/Δ^*;Tet2*^Δ/Δ^ mice suggested that severe pathology was not caused by an inability to control the pathogen, but by impaired disease tolerance. In line with this idea, *Slc23a2*^Δ/Δ^*;Tet2*^Δ/Δ^ mice typically became moribund after the day 7 parasitemia peak (**Figure 6H**). B cells and T cells help clear blood stage *Plasmodium*^51,52^. Infected *Slc23a2*^Δ/Δ^*;Tet2*^Δ/Δ^ mice had reduced B cell frequency in the marrow and changes in T cell frequence in the spleen and blood compared to infected controls (**Figure S6C-D**) but were mostly able to control blood stage *P. chabaudi* (**Figure 5E**) suggesting impaired adaptive immunity was not the major reason for increased pathology. If *Slc23a2;Tet2* deficiency promotes mortality by altering myeloerythroid hematopoiesis rather than by impairing antimicrobial immunity, then it could also impair resistance to sterile inflammation. To test that we challenged mice with a lipopolysaccharide (LPS)-induced sepsis model. Sepsis is characterized by an uncontrolled inflammatory response to infection that culminates in organ failure and death^53^. *Slc23a2^Δ/Δ^;Tet2^Δ/Δ^* mice had markedly increased mortality after LPS administration (**Figure 6J**). Together, these data suggest that ascorbate and TET2 activity in hematopoietic cells is required for tolerance to infectious or inflammatory challenge.

### Hematopoietic *Tet2;Tet3* deficiency impairs tolerance to malaria

Because *Tet2;Tet3* deletion had similar effects as *Slc23a2;Tet2* deletion on myelopoiesis (**Figure 4**), we tested if it similarly compromised balanced myeloerythroid hematopoiesis and morbidity in malaria. Single *Tet2* deletion caused minor changes in marrow myeloid cells, and minor or no changes in erythropoiesis, anemia, and body weight in infection, consistent with the lack of myeloid skewing (**Figure S7A-F**). Similar to *Slc23a2;Tet2* deletion*, Tet2*;*Tet3* deletion increased the percentage but not the number of infected erythrocytes (**Figure 6K-L**). Similar to *Slc23a2;Tet2* deletion*, Tet2*;*Tet3* deletion elevated inflammatory monocytes and neutrophils in the marrow, blood and spleen at baseline, which persisted after infection, and further increased them in the liver and lung, suggesting high myelopoiesis and tissue inflammation (**Figure 6M**). *Tet2*;*Tet3* deletion blocked the infection-induced production of bone marrow CFU-E and CD71^+^Ter119^+^ erythroblasts and exacerbated malarial anemia (**Figure 6N-T, Figure S7G-L**). *Tet2*;*Tet3* deletion exacerbated malarial hypoglycemia but did not elevate liver enzymes compared to infected control mice (**Figure 6U, Figure S7M**). Therefore, combined *Tet2;Tet3* deletion recapitulates imbalanced myeloerythroid hematopoiesis, and some but not all aspects of impaired disease tolerance caused by *Slc23a2;Tet2* deletion.

## DISCUSSION

Despite strong public interest in the use of ascorbate (vitamin C) in infections, two challenges have limited understanding of its role, if any. It is normally abundant in organisms, and supplementation when baseline levels are normal or high only marginally changes its levels and cannot test physiological roles. To identify physiological roles, depletion experiments are required^54^. And ascorbate had not been manipulated in a cell-type specific manner in infection models, obscuring its mechanism of action. We used *Slc23a2* conditional knockouts to show that naturally occurring ascorbate acts in oligopotent and restricted myeloid progenitors via TET2/TET3 to restrain myelopoiesis. Ascorbate depletion in *Gulo^−/−^*mice regulates TET activity, HSC function, and leukemogenesis similarly to *Slc23a2* deletion^5,55^, suggesting effects of *Slc23a2* deletion are mediated by ascorbate depletion. *Slc23a2* deletion does not deplete plasma ascorbate^6^ and erythrocytes use other transporters to import ascorbate^56^, suggesting that ascorbate modulates malaria pathology through its action in hematopoietic cells rather than parasitized erythrocytes. Our results localize the action of a metabolite to a defined progenitor cell and suggest a mechanism by which ascorbate ameliorates the severity of inflammation or infection.

Infections stimulate myelopoiesis and impair erythropoiesis to cause inflammation and anemia^57,58^. Inflammatory cytokines drive this effect^59–61^. In contrast the mechanisms that restrain myelopoiesis to maintain a balanced myeloerythroid output in infections are less well understood. Our results suggest this requires intact ascorbate-TET activity. *Tet2* loss in mice promotes myelopoiesis and inflammation^37,39,62–64^. *TET2-*mutant human HSPCs have increased myeloid and impaired erythroid differentiation in culture conditions that typically lack ascorbate^65–67^. The molecular mechanisms by which TET activity regulates myeloid or erythroid differentiation include hydroxymethylation of gene regulatory regions for expression of transcription factors and other lineage-specific genes^65,68–70^. Somatic *TET2* loss in hematopoietic stem and progenitor cells is common in humans and promotes clonal hematopoiesis, clonal cytopenias, and myeloid neoplasms^71^. Both *TET2-*mutant clonal hematopoiesis^72^ and ascorbate depletion^16–20^ are associated with incidence or severity of infections. Our results predict that maintaining high ascorbate levels in individuals carrying myeloid-driver mutations can preserve balanced hematopoiesis and ameliorate inflammatory pathology. This finding may provide one explanation for recent results that ascorbate supplementation of patients with clonal cytopenias or early-stage myeloid neoplasms improves survival in part by reducing mortality from severe infections^28^.

Malaria afflicts 280 million people annually, and in 2024 killed 610,000 people^73^. A conspicuous feature of malaria is that high parasitemia does not necessarily cause severe disease indicating pathogen tolerance^3^. Tolerance is promoted by pathogen exposure that changes monocyte and T cell function to attenuate inflammation^74–76^. It remains unclear to what extent tolerance requires regulation of hematopoietic stem or progenitor cells. A major cause of malarial mortality is anemia due to hemolysis and dysregulated erythropoiesis^77,78^. Inflammatory myeloid cells also contribute to pathology^44,79^. Our results suggest that a balanced hematopoietic output that maintains erythropoiesis and attenuates myelopoiesis is required for malaria tolerance. Mice with ascorbate and TET2 depletion in hematopoietic cells remain able to contain the pathogen, yet succumb to a normally non-lethal malaria infection. Ascorbate depletion is prevalent in malaria because ascorbate levels are low in the populations of malaria-endemic regions compared to the U.S.^80^ and are further reduced in malaria^47^. Somatic *TET2* mutations are prevalent in old age, raising the question of whether they are relevant to severe human malaria, which is more prevalent in children. The pediatric prevalence of severe malaria in endemic areas is due to the fact that children lack immunity acquired from repeated exposure. Among nonimmune individuals, old age dramatically increases the risk of severe malaria^81,82^. Even in endemic countries, the malarial mortality curve is U-shaped, with high rates in early childhood and again in late middle or old age^83,84^. Our results thus raise the possibility that variation in non-heritable host factors such as ascorbate depletion and somatic *TET2* mutations contributes to variation in malarial morbidity and mortality.

## METHODS

### Mice

*Slc23a2^fl/fl^* ^6^, *Tet2^fl/fl^* (IMSR_JAX:017573)^37^, *Tet3^fl/fl^* (IMSR_JAX:031015)^85^, *Mx1-cre* (IMSR_JAX:003556)^86^, and *Vav1-cre* (IMSR_JAX:008610)^87^ mice were described previously. All mice were on a C57BL background. Both male and female mice were used in all studies. Mice were injected with 40 μg poly (I:C) (Cytiva) every other day for a total of five intraperitoneal injections starting at 4-6 weeks of age to induce Cre expression. Mice were analyzed 4-6 weeks after the first day of poly (I:C) injection, or as indicated in the text. For sepsis induction, mice received an intraperitoneal injection of 2.5 mg/kg LPS (Sigma Aldrich). C57BL/Ka-Thy-1.1 (CD45.2) and C57BL/Ka-Thy-1.2 (CD45.1) mice were used for transplantation experiments. Mice were housed in the Animal Resource Center of UT Southwestern and all procedures were approved by the UT Southwestern Institutional Animal Care and Use Committee.

### Hematopoietic cell isolation and hematopoietic analysis

Bone marrow cells were obtained by flushing femurs and tibia with a 25G needle, or crushing femurs, tibias, vertebrae, and pelvic bones with a mortar and pestle in staining medium consisting of Ca^2+^/Mg^2+^-free Hank’s balanced salt solution (HBSS; Gibco), supplemented with 2% heat-inactivated bovine serum (Gibco). Spleens were mechanically dissociated by trituration in staining medium. Livers and lungs were enzymatically digested for 30 minutes at 37°C in 1.5 ml RPMI-1640 (Sigma-Aldrich), containing 250 μg/ml liberase (Roche) and 100 μg/ml DNase I (Roche). Cell suspensions were filtered through a 40 μm strainer. Cell number was measured with a Vi-CELL cell viability analyzer (Beckman Coulter). Blood was collected by cardiac puncture using a 25G needle and mixed in a tube containing 5 μl 0.5 M EDTA. Complete blood cell counts were determined using a hemavet HV950 (Drew Scientific) or Element HT5 (Heska). For hematopoietic analysis, 40 μl blood was lysed in 1 ml of ammonium chloride buffer (155 mM NH4Cl; 10 mM KHCO3; 0.1 mM EDTA) for 10-15 minutes at 4°C. Cells were incubated with fluorescently conjugated antibodies for 90 minutes on ice when using CD34 antibody or for 30 minutes at 4°C. Cells were washed with staining media and resuspended in staining media containing 1 μg/ml DAPI or 1 μg/ml propidium iodide for live/dead discrimination. Cell populations were defined as previously described^31,33,88–90^ with the following markers: CD150^+^CD48^−^Lineage^−^Sca-1^+^Kit^+^ hematopoietic stem cells (HSCs), CD150^−^CD48^−^Lineage^−^Sca-1^+^Kit^+^ multipotent progenitor cells (MPPs), CD150^−^CD48^+^Lineage^−^Sca-1^+^Kit^+^ hematopoietic progenitor cells (HPC-1), CD150^+^CD48^+^Lineage^−^Sca-1^+^Kit^+^ hematopoietic progenitor cells (HPC-2), CD34^+^CD16/32^−^Lineage^−^Sca-1^−^Kit^+^ common myeloid progenitors (CMPs), CD34^+^CD16/32^+^Lineage^−^Sca-1^−^Kit^+^ granulocyte-monocyte progenitors (GMPs), CD34^−^CD16/32^−^Lineage^−^Sca-1^−^Kit^+^ megakaryocyte-erythroid progenitors (MEPs), Ly6C^−^CD115^−^CD34^+^CD16/32^+^Lineage^−^Sca-1^−^Kit^+^ granulocyte-monocyte progenitors (Ly6C^−^CD115^−^GMPs), CD150^+^CD105^−^CD41^−^CD16/32^−^Lineage^−^Sca-1^−^Kit^+^ pre-megakaryocyte-erythrocyte progenitors (Pre-MegE), CD150^−^CD105^−^CD41^−^CD16/32^−^Lineage^−^Sca-1^−^Kit^+^ pre-granulocyte-macrophage progenitor (Pre-GM), CD150^+^CD41^+^Lineage^−^Sca-1^−^Kit^+^ megakaryocyte progenitors (MkP), CD150^+^CD105^+^CD41^−^CD16/32^−^Lineage^−^Sca-1^−^Kit^+^ pre-colony-forming-unit-erythroid progenitors (Pre-CFU-E), CD150^−^CD105^+^CD41^−^CD16/32^−^Lineage^−^Sca-1^−^Kit^+^ colony-forming-unit-erythroid progenitors (CFU-Es), Mac-1^+^ myeloid cells, Mac1^+^CD115^+^Ly6C^+^Ly6G^−^inflammatory monocytes, Mac1^+^CD115^−^Ly6C^mid^Ly6G^+^ neutrophils, Mac1^−^B220^+^ B cells, Mac1^−^CD3^+^ T cells, Mac1^−^B220^−^CD3^−^CD71^mid^Ter119^−^immature erythroid progenitors, Mac1^−^B220^−^CD3^−^CD71^+^Ter119^−^erythroid progenitors, Mac1^−^B220^−^CD3^−^CD71^+^Ter119^+^ erythroblasts. The Lineage cocktail for HSCs and progenitors consisted of CD2, CD3, CD5, CD8, Ter119, B220, and Gr-1 antibodies. CD86 was used instead of Sca-1 to define HSPCs in experiments that measured 5hmC levels in malaria infection. For all flow cytometry experiments, combinations of antibodies were used against the following cell-surface markers, conjugated to Violet Fluor 450, Brilliant Violet 421, Brilliant Violet 510, FITC, PE, PerCP-Cy5.5, PE-Cy7, APC, eFluor 660, Alexa Fluor 700, APC/Fire 750, or APC-Cy7: CD2 (RM2-5, Tonbo Biosciences/BioLegend), CD3 (17A2, Tonbo Biosciences/BioLegend), CD5 (53-7.3, BioLegend), CD8a (53-6.7, Tonbo Biosciences/BioLegend), CD11b (Mac1) (M1/70, Tonbo Biosciences/BioLegend), CD16/32 (93, BioLegend), CD34 (RAM34, Invitrogen), CD41 (MWReg30, BioLegend), CD45.1 (A20, Tonbo Biosciences), CD45.2 (104, Tonbo Biosciences/BioLegend), CD45R (B220) (RA.3-6B2, Tonbo Biosciences/BioLegend), CD48 (HM48-1, BioLegend), CD71 (R17217, BioLegend), CD86 (GL-1, BioLegend), CD105 (MJ7/18, BioLegend), CD115 (CSF-1R) (AFS98, BioLegend), CD117 (c-Kit) (2B8, BioLegend), CD150 (TC15-12F12.2, BioLegend), Ly-6A/E (Sca-1) (D7, BioLegend), Ly-6C (HK1.4, BioLegend), Ly-6G (Gr-1) (RB6-8C5, Tonbo Biosciences; 1A8, BioLegend), or Ter119 (TER119, Tonbo Biosciences/BioLegend). Analysis and sorting were performed using the FACSAria flow cytometer (BD Biosciences), FACSymphony (BD Biosciences) or a FACSCanto (BD Biosciences). Data were analyzed using FlowJo (Flowjo LLC) or FACSDiva (BD Biosciences).

### Hematopoietic progenitor transplantation assays

Recipient mice (CD45.1) were lethally irradiated using an XRAD 320 X-ray irradiator (Precision X-Ray) with two doses of 5.4 Gy (10.8 Gy total) delivered at least 3 hours apart. For HPC-1 transplants, we transplanted 1,000 purified CD150^−^CD48^+^Lin^−^Sca1^+^Kit^+^ HPC-1 cells (CD45.2) with 400,000 wild-type bone marrow competitor cells (CD45.1/CD45.2) into the retro-orbital venous sinus of anesthetized recipient (CD45.1) mice. For CMP transplants, we transplanted 5,000 purified CD34^+^CD16/32^−^Lin^−^Sca1^−^Kit^+^ CMP cells (CD45.2) with 400,000 wild-type bone marrow competitor cells into anesthetized recipient (CD45.1) mice. Recipient mice were maintained on antibiotic water (Baytril 0.08 mg/ml) for 1 week pre-transplantation and during the transplant. Bone marrow cells for analysis of transplant recipient mice were obtained by crushing femurs, tibias, vertebrae, and pelvic bones.

### 5hmdC, 5mdC and dC measurement by LC-MS/MS

To measure modified cytosines in the DNA of purified populations, cells were sorted by flow cytometry into Ca^2+^/Mg^2+^-free HBSS supplemented with 2% heat-inactivated bovine serum, washed twice with ice-cold PBS, and the genomic DNA was extracted using Quick-DNA Miniprep kit (Zymo Research) according to the manufacturer’s instructions with the following modifications. Lysis buffer was supplemented with 2.4U/sample of proteinase K (New England Biolabs) and 0.2 mg/ml RNase A (New England Biolabs) and samples were eluted with 2 x 11 μl H2O containing 40 μM butylated hydroxytoluene (BHT, Sigma-Aldrich). Total genomic DNA from sorted cells was digested for 3 hours at 37°C with 7.5U/sample DNA Degradase Plus (Zymo Research), 10U/sample micrococcal nuclease (Thermo Scientific) and the following internal standards were added to the mix: ^13^C,^15^N2-2-deoxycytidine (^13^C,^15^N2-^2^dC), D3-5-methyl-2-deoxycytidine (D3-5mdC), D3-5-hydroxymethyl-2-deoxycytidine (D3-5hmdC) (Toronto Research Chemicals). Samples were then filtered with a Nanosep 3k column to remove enzymes, and filtrates were analyzed by LC–MS/MS. For 5mC and 5hmC, liquid chromatography was carried out with a Scherzo SM-C18 (Imtakt) reverse phase C18 column, 3 μm, 2.1× 100 mm coupled to an AB Sciex QTRAP 6500+ or 7500 operating in MRM mode as previously described^13^. Chromatographic separation was performed using a flow rate of 0.5 mL/min. The mobile phases consisted of LC-MS-grade water containing 0.1% (v/v) formic acid (mobile phase A) and LC-MS-grade acetonitrile containing 0.1% (v/v) formic acid (mobile phase B). The gradient started at 0% B from 0–2.5 min, followed by a linear increase to 100% B from 2.5–24 min. The gradient was maintained at 100% B until 25 min, then decreased to 0% B at 27 min and maintained at 0% B until the end of the run at 30 min. Data analysis was performed with MultiQuant (SCIEX).

### *Plasmodium chabaudi* infection and parasitemia measurements

*Plasmodium chabaudi* AS was originally provided by J. Langhorne, Francis Crick Institute, to the Takahashi laboratory. Malaria infections in mice were performed as previously described^91^, by intraperitoneal injection of 1×10^6^ *P. chabaudi*-infected red blood cells in 100 μl. Infected red blood cells were obtained by collecting blood by cardiac puncture from donor mice infected with 1×10^7^ cryopreserved *P. chabaudi*-infected red blood cells. Parasitemia was monitored either by microscopic counting of Giemsa-stained blood smears or by flow cytometry. For parasitemia measurements by flow cytometry, collected blood was diluted 1:10 with HBSS containing 10 mM EDTA and 2 μl blood was stained for 30 minutes at RT in the dark in 200 μl 1% BSA/PBS with the following antibodies: Ter119-FITC (Biolegend, TER-119, 1/200), CD71-PE (Biolegend, RI7217, 1/200), and Hoechst 33342 (Invitrogen, H3570, 1/1000). Analysis was performed using a FACSAria flow cytometer (BD Biosciences). The % of *P. chabaudi*-infected red blood cells was calculated as the % of CD71^−^Ter119^+^ cells that were Hoechst^+^ in infected mice over an uninfected mouse control.

### Liver damage and glucose measurements

Blood was collected by cardiac puncture using a 25G needle, mixed in a tube containing 10 μl 0.5M EDTA and centrifuged at 6,000 rpm for 2 minutes. Plasma (supernatant) was transferred to a new tube, flash-frozen in liquid nitrogen and stored at −80°C until analysis. Alanine aminotransferase (ALT), aspartate aminotransferase (AST), and glucose levels were quantified by the UT Southwestern Metabolic Phenotyping Core using VITROS MicroSlide™ technology.

### Statistical analysis

Figure panels show the pooled results from mice analyzed from multiple independent experiments. Prior to analyzing the statistical significance of differences among treatments, we tested whether data were normally distributed with the D’Agostino-Pearson test or, when n < 8, the Shapiro–Wilk test, and whether variability significantly differed between treatments with F-tests. If the data did not significantly deviate from normality, we used a parametric test and if the data significantly deviated from normality (p < 0.01 for at least one treatment), a non-parametric test. To assess the statistical significance of differences in means between treatments, we used a *t*-test or one-way ANOVA for data that was normally distributed and had equal variability, or a *t*-test with Welch’s correction or Brown-Forsythe ANOVA for data that was normally distributed and had unequal variability, or a Mann-Whitney or Kruskal-Wallis test for data that was not normally distributed. Results were graphed and statistical analysis performed using GraphPad Prism v11.

## ACKNOWLEDGEMENTS

The work was funded by grants from the Cancer Prevention and Research Institute of Texas (CPRIT) (RR180007, RP250280, RP260427), American Society of Hematology (Faculty Scholar award), Moody Foundation, an American Cancer Society and Simmons Cancer Center Institutional Research grant (IRG-17-174-13, P30CA142543), the Haggerty Foundation, and the National Institutes of Health (R01DK125713, R01HL161387) to MA. FRF is supported by a New Innovator Award by the National Institutes of Health (DP2GM154010) and a Howard Hughes Medical Institute Freeman Hrabowski Scholar award. We thank the North Texas Clinical Pharmacology Cancer Core (supported by the CPRIT Core Facilities Support Award RP210209 to LL) for 5hmC analysis, Michael Ortiz and the CRI Flow cytometry facility for flow cytometry support, the UT Southwestern Metabolic Phenotyping Core for quantitation of ALT, AST, and glucose levels, Dieu Linh Nguyen, Qing Ding, and Shiu Jie Ling for mouse colony management and technical assistance, Sean Morrison and Stefano Comazzetto for sharing *Slc23a2^fl^* mice, and Anjana Rao and Ageliki Tsagaratou for sharing *Tet3^fl^* mice.

---

**Supplemental Figure 1.**
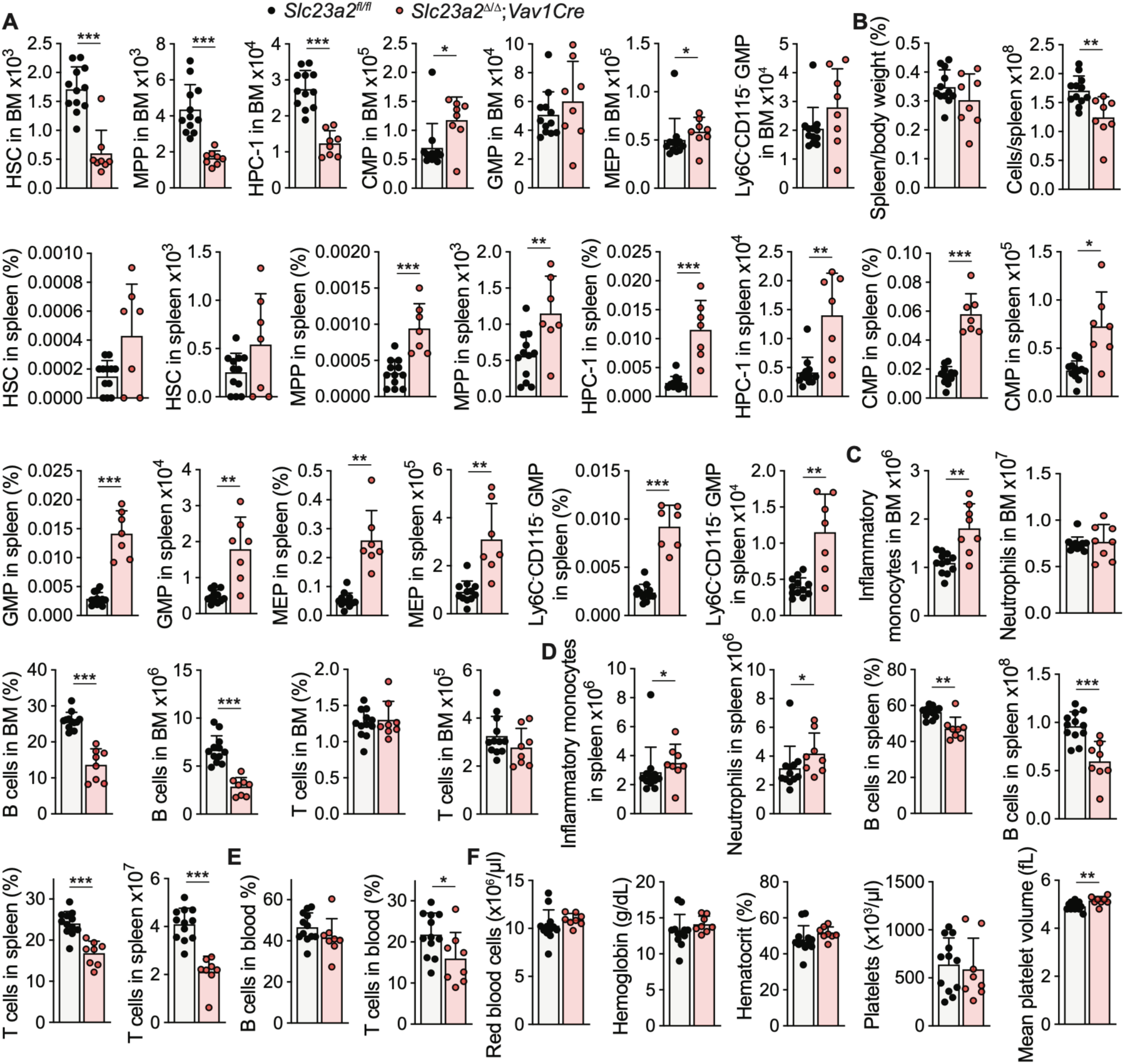
Myeloid skewing in *Vav1Cre;Slc23a2*^Δ/Δ^ mice. (**A**) Numbers of HSCs and progenitors in the bone marrow of *Vav1Cre;Slc23a2*^Δ/Δ^ and littermate control mice at 8 weeks of age. (**B**) Relative spleen size, spleen cellularity, HSPC frequencies and cell numbers in the spleen. (**C-F**) Frequencies and numbers of mature cells in the bone marrow, spleen, and blood, and blood cell counts of *Vav1Cre;Slc23a2*^Δ/Δ^ and littermate control mice.

**Supplemental Figure 2.**
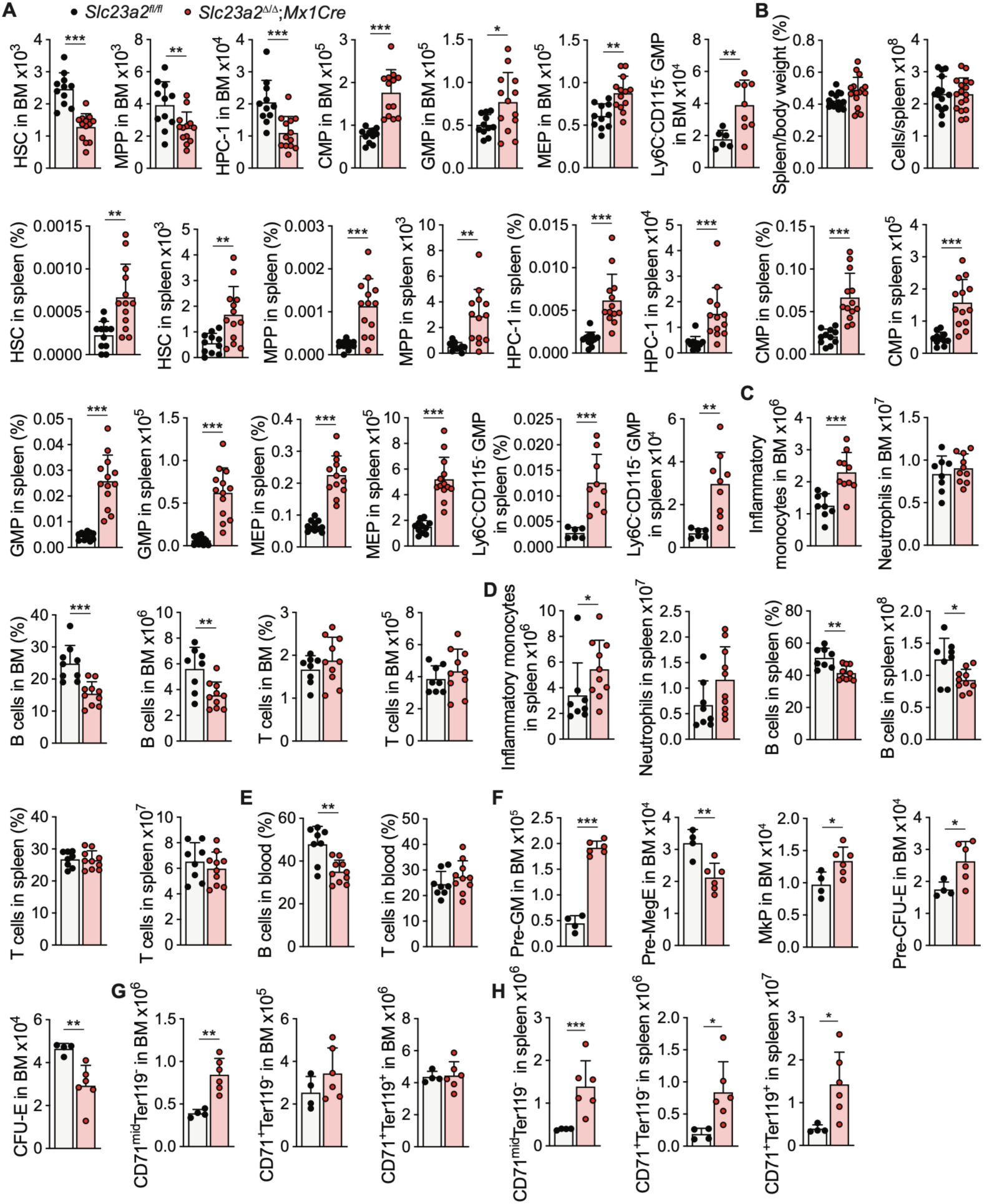
Myeloid skewing in *Mx1Cre;Slc23a2*^Δ/Δ^ mice. (**A**) Numbers of HSCs and progenitors in the bone marrow of *Mx1Cre;Slc23a2*^Δ/Δ^ and littermate control mice at 4-8 weeks after *Mx1Cre* induction with poly(I:C). (**B**) Relative spleen size, spleen cellularity, HSPC frequencies and cell numbers in the spleen. (**C-E**) Frequencies and cell numbers of mature cells in the bone marrow, spleen, and blood. (**F**) Numbers of early myeloid, megakaryocyte, and erythroid progenitors in the bone marrow. (**G-H**) Numbers of late-stage erythroid populations in the bone marrow and spleen.

**Supplemental Figure 3.**
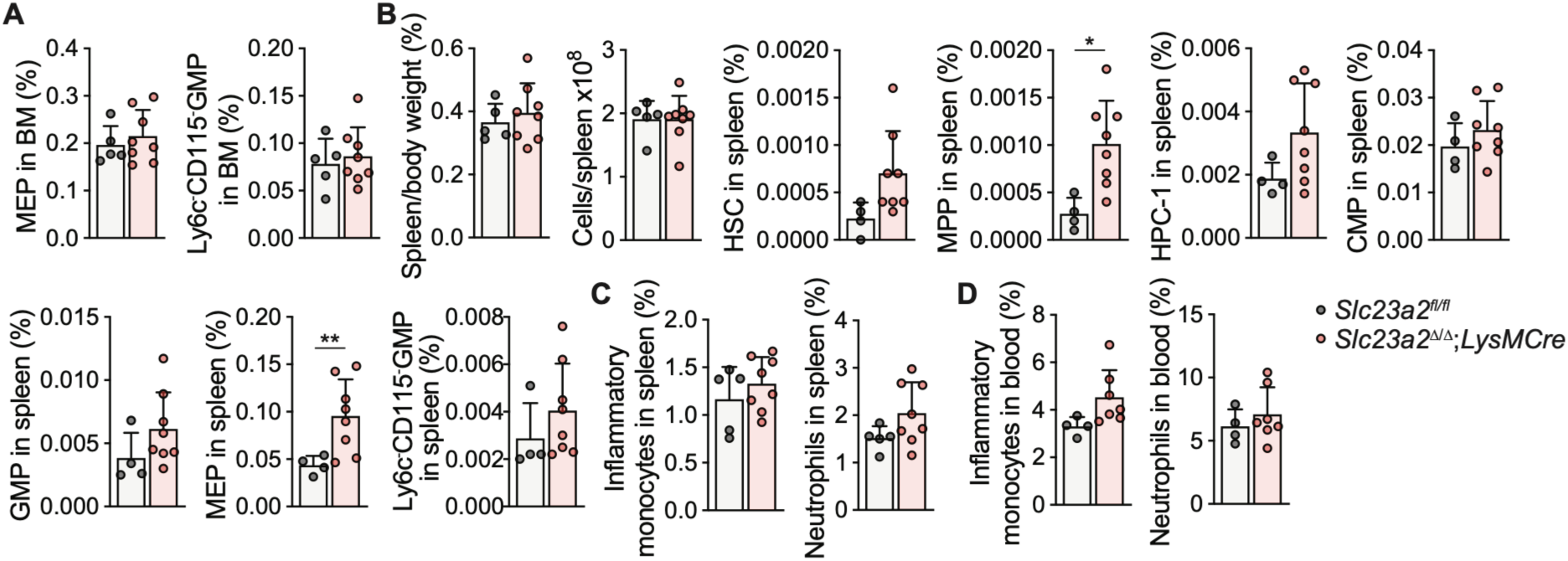
Ascorbate deficiency in mature myeloid cells does not affect myelopoiesis. (**A**) Frequencies of myeloid progenitors in the bone marrow of wild-type and *LysMCre;Slc23a2*^Δ/Δ^ mice at 8 weeks of age. (**B**) Spleen size, spleen cellularity and frequencies of HSCs and progenitors in the spleen. (**C-D**) Frequencies of mature cells in the spleen and blood.

**Supplemental Figure 4.**
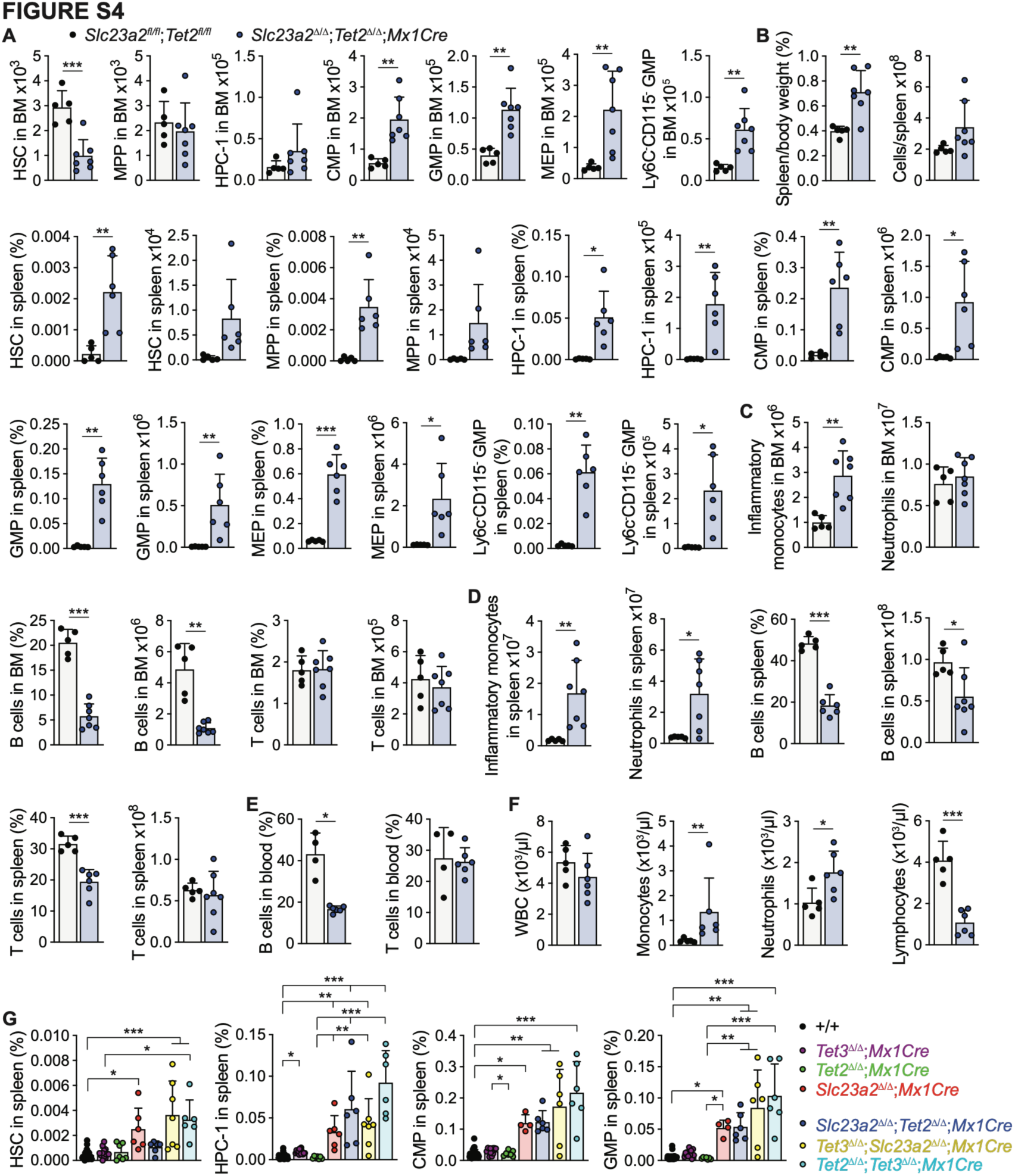
T*e*t2 deficiency collaborates with ascorbate deficiency to increase myeloid skewing. (**A**) Numbers of HSCs and progenitors in the bone marrow of *Mx1Cre;Slc23a2*^Δ/Δ^*;Tet2*^Δ/Δ^ and littermate control mice at 4-8 weeks after deletion. (**B**) Relative spleen size, spleen cellularity, HSPC frequencies and numbers in the spleen. (**C-F**) Frequencies and numbers of mature cells in the bone marrow, spleen, and blood, and blood cell counts of *Mx1Cre;Slc23a2*^Δ/Δ^*;Tet2*^Δ/Δ^ and littermate control mice. (**G**) Frequencies of HSCs and progenitors in the spleen of mice from the indicated genotypes. +/+ refers to littermate control mice carrying the corresponding floxed alleles.

**Supplemental Figure 5.**
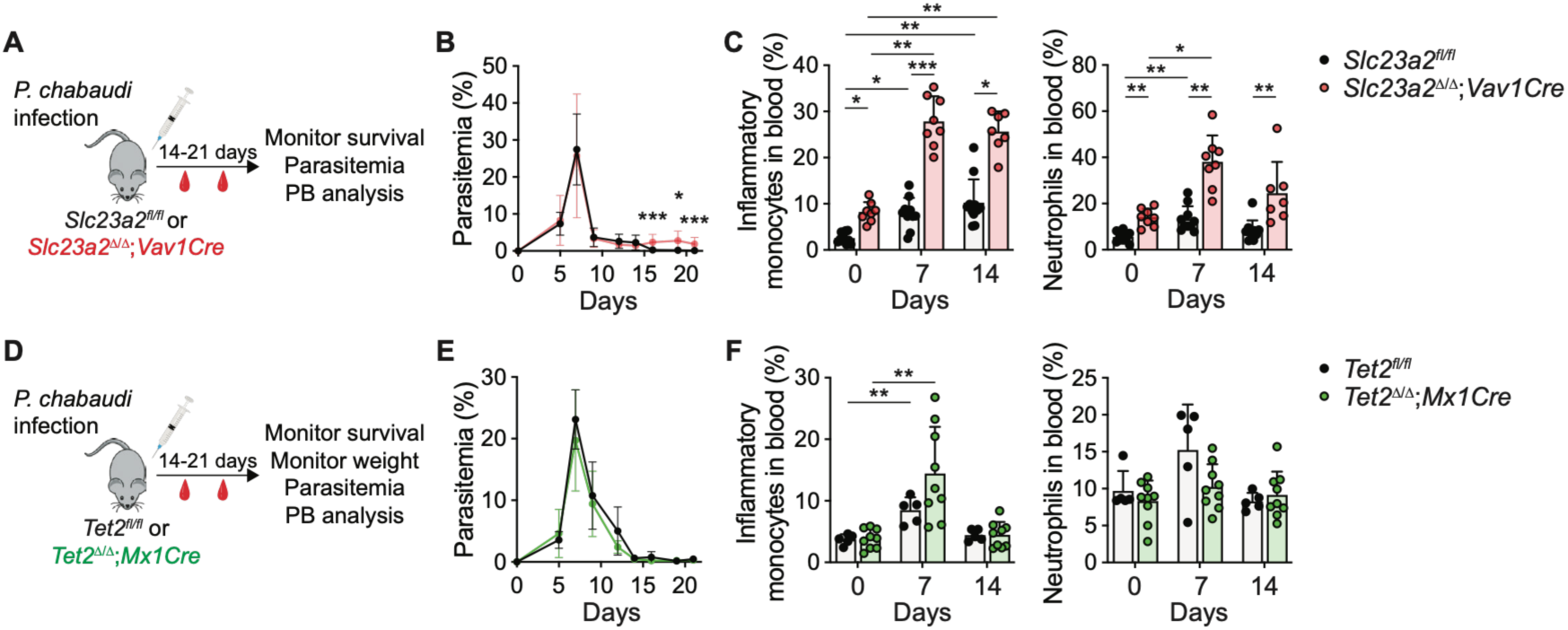
Ascorbate deficiency alone but not TET2 deficiency alone increases the myeloid response to *Plasmodium chabaudi* infection. Parasitemia and peripheral blood analysis in *P. chabaudi* infected *Vav1Cre;Slc23a2*^Δ/Δ^ vs littermate control mice (**A-C**) or *Mx1Cre;Tet2*^Δ/Δ^ vs littermate control mice (**D-F**).

**Supplemental Figure 6.**
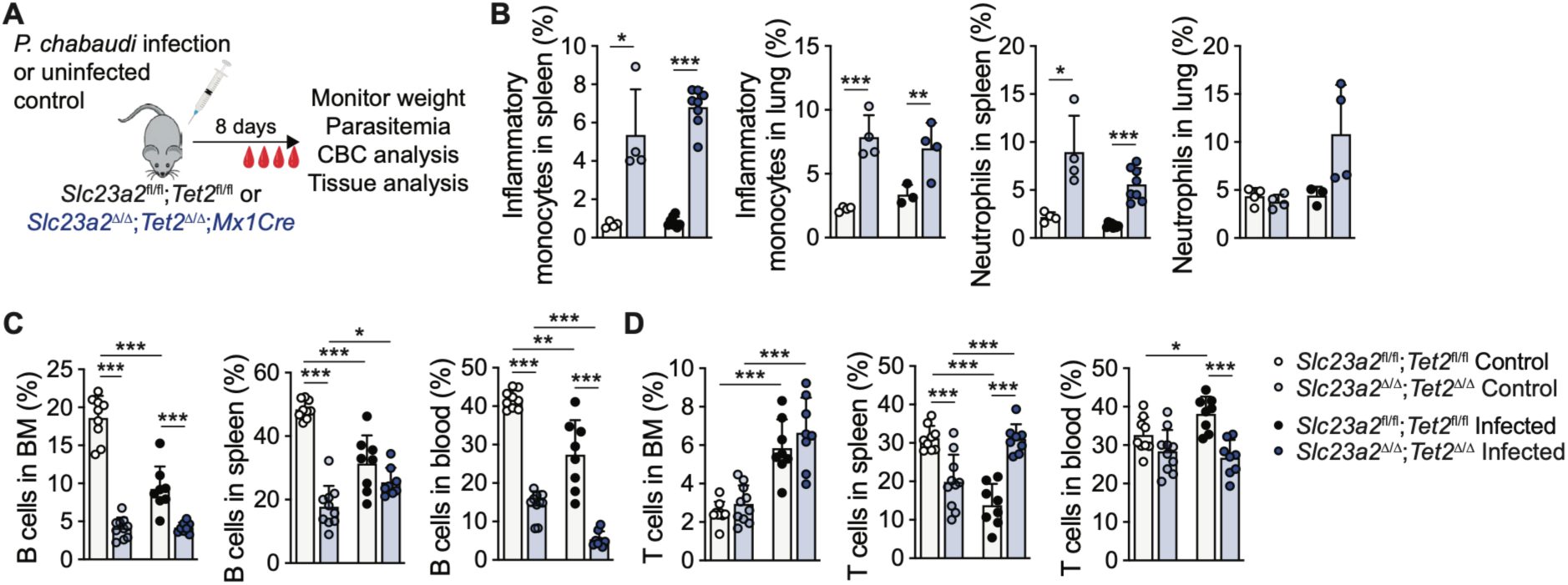
Effects of combined *Slc23a2* and *Tet2* deletion on malarial hematopoiesis. Analysis of myeloid and lymphoid cells in the bone marrow, spleen, blood, and lung in uninfected or *P. chabaudi*-infected mice.

**Supplemental Figure 7.**
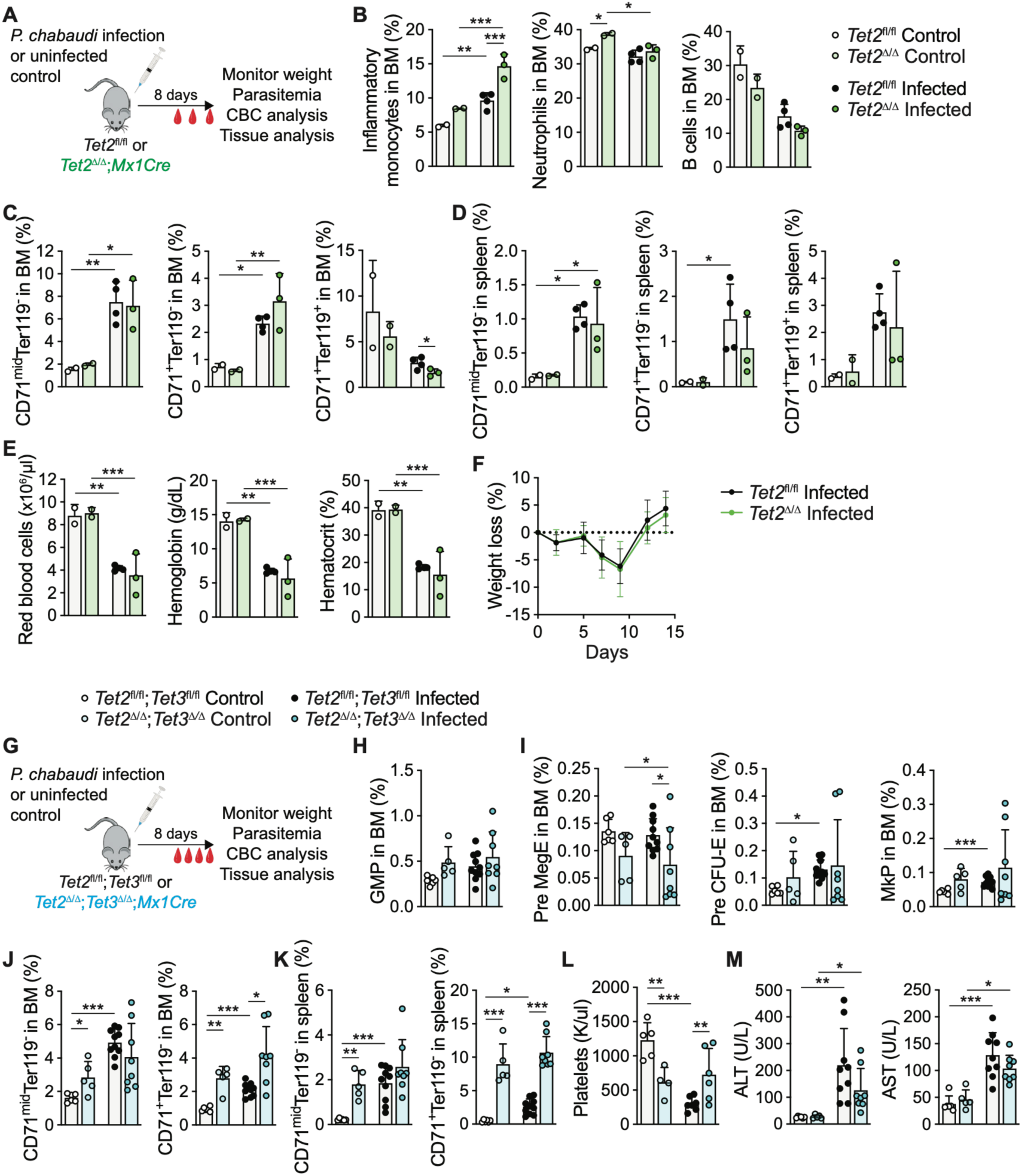
Effects of *Tet2* deficiency or *Tet2;Tet3* deficiency on hematopoiesis and pathology in *P. chabaudi* infected mice. Hematopoietic analysis and malarial pathology 8 days after *P. chabaudi*-infection in wild-type vs *Mx1Cre;Tet2*^Δ/Δ^ mice (**A-F**), or wild-type vs *Mx1Cre;Tet2*^Δ/Δ^*;Tet3*^Δ/Δ^ mice (**G-M**).

